# Adaptive optical two-photon fluorescence microscopy probes cellular organization of ocular lenses in vivo

**DOI:** 10.1101/2023.01.17.524320

**Authors:** Santosh Kumar Paidi, Qinrong Zhang, Yuhan Yang, Chun-Hong Xia, Na Ji, Xiaohua Gong

## Abstract

The mammalian ocular lens is an avascular multicellular organ that grows continuously throughout life. Traditionally, its cellular organization is investigated using dissected lenses, which eliminates in vivo environmental and structural support. Here, we demonstrated that two-photon fluorescence microscopy (2PFM) can visualize lens cells in vivo. To maintain subcellular resolution at depth, we employed adaptive optics (AO) to correct aberrations due to ocular and lens tissues, which led to substantial signal and resolution improvements. Imaging lens cells up to 980 μm deep, we observed novel cellular organizations including suture-associated voids, enlarged vacuoles, and large cavities, contrary to the conventional view of a highly ordered organization. We tracked these features longitudinally over weeks and observed the incorporation of new cells during growth. Taken together, non-invasive longitudinal in vivo imaging of lens morphology using AO 2PFM will allow us to directly observe the development or alterations of lens cellular organization in living animals.

## Introduction

The ocular lens, together with the cornea, is responsible for focusing the light on the retina. Composed of fiber cells and an anterior monolayer of epithelial cells, the lens continuously grows throughout life, with new fiber cells differentiating from the equatorial epithelial cells and incorporated at the periphery of the lens^1–3^. Lacking vasculature, the lens maintains its transparency by the loss of organelles in the maturing fiber cells^4,5^ and engages in active microcirculation to facilitate the transport of ions and small molecules^3,6,7^. Various diseases affect the ocular lens by causing structural disruptions and physiological changes on the cellular level.^1,8–10^ For example, presbyopia and age-related cataracts involve chronic changes in the interior fiber cells, whose cellular mechanisms remain poorly understood^11–19^. To unravel the biological underpinnings of physiological and pathological transformations in the lens, we need to study cellular changes longitudinally in vivo with minimal invasiveness.

Due to the strong refractive power of the ocular lens, in vivo imaging of its internal structure using standard microscopy methods is hampered by the substantial optical aberrations that lens cells introduce into the image-formation process. Likely as a result, current knowledge about the cellular organization of the lens is mainly based on ex vivo light and electron microscopy studies of enucleated mouse and human lenses^2,20–27^, which exclude the possibility of longitudinal investigations. Furthermore, an enucleated lens lacks its in vivo environment such as the lens internal circulation current and attached lens zonular structures^28^. Sample processing steps such as vibratome sectioning, fixation, and staining can also disrupt cellular organization. Therefore, there is an unmet need for an imaging tool that can non-invasively and longitudinally monitor cells deep inside the ocular lens at subcellular resolution.

Here we applied two-photon fluorescence microscopy (2PFM)^29^, a technique widely used to image brain tissues^30^, to in vivo imaging of the mouse anterior lens up to ~1 mm deep under the epithelium. Using adaptive optics (AO), we corrected the wavefront aberrations introduced by lens cells, which led to substantial improvements in signal and resolution and enabled subcellular resolution imaging up to 700 μm into the lens interior. By employing wildtype mice expressing tdTomato^31^ in the plasma membrane of lens cells^26^ and variants generated by crossing them with transgenic mice^21^ with homozygous and heterozygous deletion of KLPH gene, we discovered hitherto unreported features such as anterior voids, enlarged vacuoles in the fiber cells, and non-uniform organization and shapes of fiber cells near the suture regions of the adult mice across all the strains. Our observations indicate that the anterior organization of fiber cell tips near the lens suture varies with depth. We repeatedly imaged the lenses of the same mice and visualized changes in cellular organization during their growth. To our knowledge, our study is the first in vivo 2PFM study of lenses as well as the first demonstration of longitudinal tracking of distinct morphological features in the live mouse eye. Capable of non-invasive longitudinal characterizations of cellular organization in the mouse lens in vivo, AO 2PFM can be generally applied to studying physiological events during lens development and pathological transformations in disease models at subcellular resolution.

## Results

### AO enables diffraction-limited 2PFM imaging of cells deep in the mouse ocular lens in vivo

In 2PFM, a near-infrared (NIR) excitation laser is tightly focused into a fluorescent sample, where two NIR photons need to be absorbed simultaneously by a fluorophore to generate a fluorescence photon. The miniscule cross sections of two-photon absorption confine fluorescence generation to the focus of the excitation laser. Therefore, 2PFM is capable of optically sectioning thick tissues and extracting information only from the focal plane of the microscope objective. Scanning the excitation focus in 3D and collecting the emitted fluorescence at each position with a non-imaging detector (e.g., a photomultiplier tube), one obtains information on fluorophore distribution and brightness. The resolving power and signal of 2PFM are determined by the spatial dimension and the intensity of the excitation laser focus, respectively. The focus that is most confined in space (leading to the highest resolving power) and has the highest focal intensity (leading to the largest fluorescence signal) is formed when the excitation light has a spherical wavefront in the vicinity of the focus. Because the ocular lens has refractive indices substantially higher than water^32^, the excitation light focused by water-dipping or water-immersion objective lenses accumulates wavefront aberrations while propagating through the ocular lens. The distorted wavefront of the excitation light leads to an enlargement of focal volume and a decrease of focal intensity, causing a degradation of both 2PFM resolution and signal^33,34^.

For in vivo lens imaging, we employed wildtype (WT, n = 6) mice expressing tdTomato^31^ in the plasma membrane of lens cells^26^ and their variants generated by crossing them with transgenic mice^21^ with homozygous (KLPH-KO, n = 5) and heterozygous (KLPH-Het, n = 4) deletion of KLPH gene. Following pupil dilation under isoflurane anesthesia, we imaged the lenses of these mice using a homebuilt 2PFM system equipped with an AO module^35^ (AO 2PFM). We imaged the anterior central part of the lens at multiple depths starting at lens epithelium using 1000 nm excitation laser. Due to the transparency of the lens, we utilized direct wavefront sensing AO to measure and correct the aberrations introduced by the lens cells. Using the tdTomato two-photon fluorescence signal itself as a wavefront-sensing guide star following previous described procedures^36,37^, we measured the eye-induced aberrations on the excitation wavefront with a Shack-Hartmann (SH) sensor and cancelled out these aberrations with a segmented deformable mirror (DM). At a depth of 160 μm below the epithelium, we imaged the lens fiber cells and found that the AO correction provided ~2x improvement in 2P fluorescence signal (Field of view: 228×228 μm^2^, **Fig. 1A-C**; 76×76 μm^2^, **Fig. 1D-F**). The corrective wavefront pattern (**Fig. 1G**) indicates a dominance of coma. The spatial frequency representations of the images show that correcting for the sample-induced aberrations resulted in a significant improvement in the lateral resolution and recovery of higher spatial frequencies up to the diffraction limit (**Fig. 1H-I**). Next, we checked to see if we can maintain subcellular resolution at greater depths below the epithelium, which was expected to be challenging due to the larger refractive power and denser fiber cell organization at the core of the lens. At a depth of 710 μm, in addition to increasing 2P fluorescence signal, AO correction became essential for resolving the fine membrane features at the anterior ends of fiber cells, which meet at the suture region (**Fig. 1J-P**). This resolution improvement was also observed in the spatial frequency representations of the images (**Fig. 1Q-R**).

**Figure 1.**
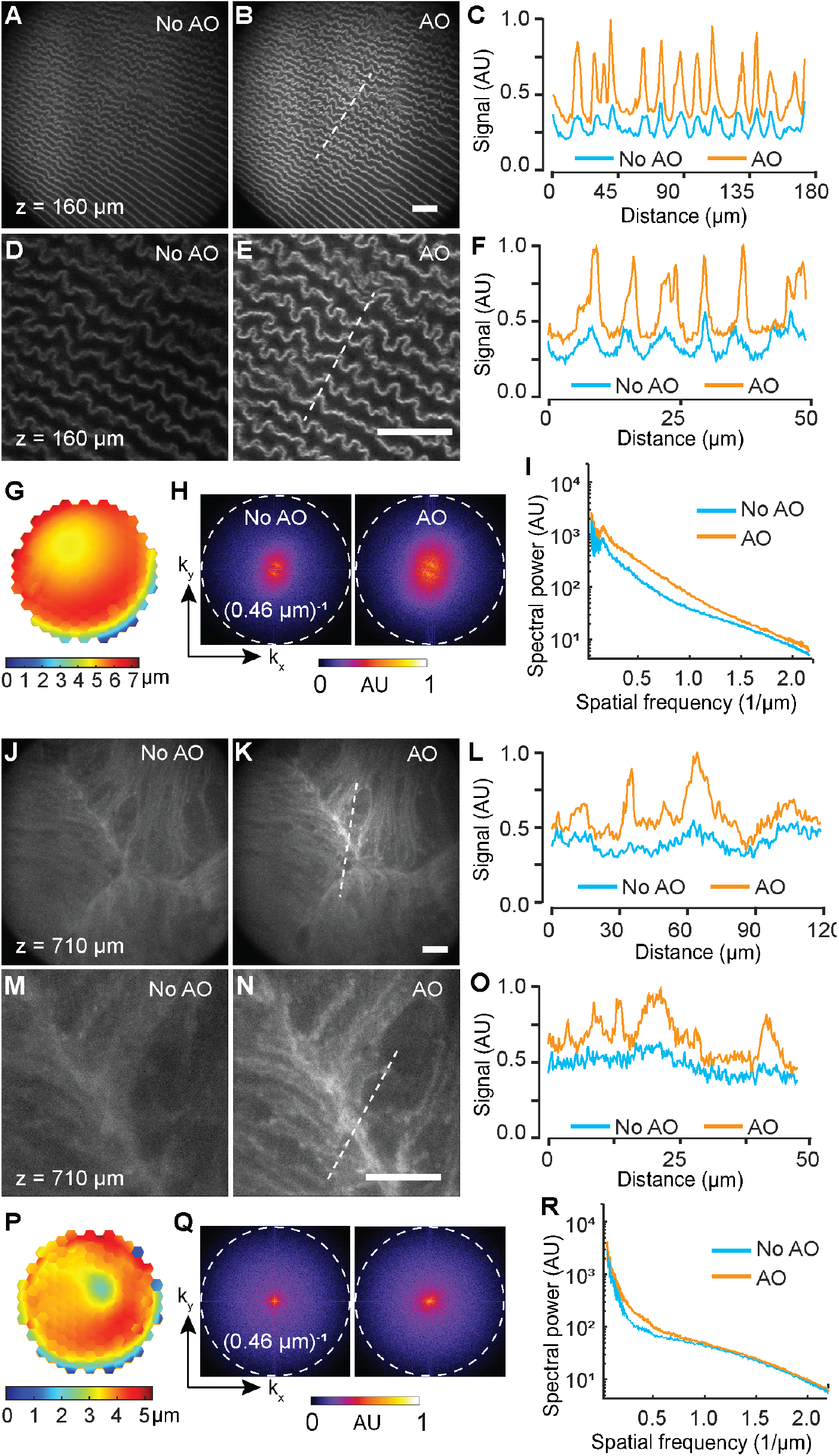
AO 2PFM enables in vivo imaging of lens cellular organization at depth. (**A-B**) 2PFM images of lens fiber cells at 160 μm below epithelium measured without and with AO, respectively. (**C**) Signal profiles along the dashed line in **A** and **B**. (**D-E**) Zoomed-in views of sub-regions from **A** and **B**. (**F**) Signal profiles along the dashed line in **D** and **E**. (**G**) Corrective wavefront for **B** and **E**. (**H-I**) Spatial frequency space representations of images in **D** and **E** and their radially averaged profiles, respectively. Dashed circles: diffraction limit at (0.46 μm)^−1^. (**J-K**) Images of lens cells at anterior suture 710 μm below lens epithelium without and with AO, respectively. (**L**) Signal profiles along the dashed line in **J** and **K**. (**M-N**) Zoomed-in views of sub-regions from **J** and **K**. (**O**) Signal profiles along the dashed line in **M** and **N**. (**P**) Corrective wavefront for **K** and **N**. (**Q-R**) Spatial frequency space representations of images in **M** and **N** and their radially averaged profiles, respectively. Scale bars: 25 μm.

While we observed consistent AO improvement of 2PFM imaging of lens cells up to ~700 μm below epithelium, correction at deeper regions of the lens was limited by the size of the dilated pupil. At these depths, the iris started to block the edge of the excitation light, leading to reduced excitation numerical aperture. The wavefront of the outgoing fluorescence signal was also clipped. Because the fluorescence wavefront was measured directly by the SH sensor to generate the corrective wavefront, such clipping also hindered the robust reconstruction of the corrective wavefront for aberration correction. Nevertheless, when we used the corrective wavefront acquired at 700 μm to correct for aberrations at deeper regions in the lens, we observed ~30% improvement in 2P signal and a substantial improvement in image resolution up to 900 μm (**Fig. S1, Supporting Information**).

### AO 2PFM reveals rich features of fiber cell organization throughout ocular lenses in live mice

We leveraged the 2P signal and resolution enhancement obtained by AO to image the organization of fiber cells at various depths (**Fig. 2**). We found similar features across the lenses of WT, KLPH-Het and KLPH-KO mouse models employed in the current study. Therefore, in the subsequent sections, we have combined the results from all the strains and summarized the strain information for individual figure panels in **Table ST1 (Supporting Information)**. Prior ex vivo studies had established an onion-like model of fiber cell organization within the lens capsule^2^. Prominent features described in the literature include a monolayer of epithelial cells that cover the anterior surface, the primary fiber cells at the core of the lens formed early during the embryonic development, and meridional rows of secondary fiber cells that are continuously incorporated into the lens at the equatorial margin of the epithelium, which elongate until their tips form anterior and posterior suture structures^1,2^. Previous studies also reported highly organized fiber cells with hexagonal cross-section and well-defined suture structures^38,39^.

**Figure 2.**
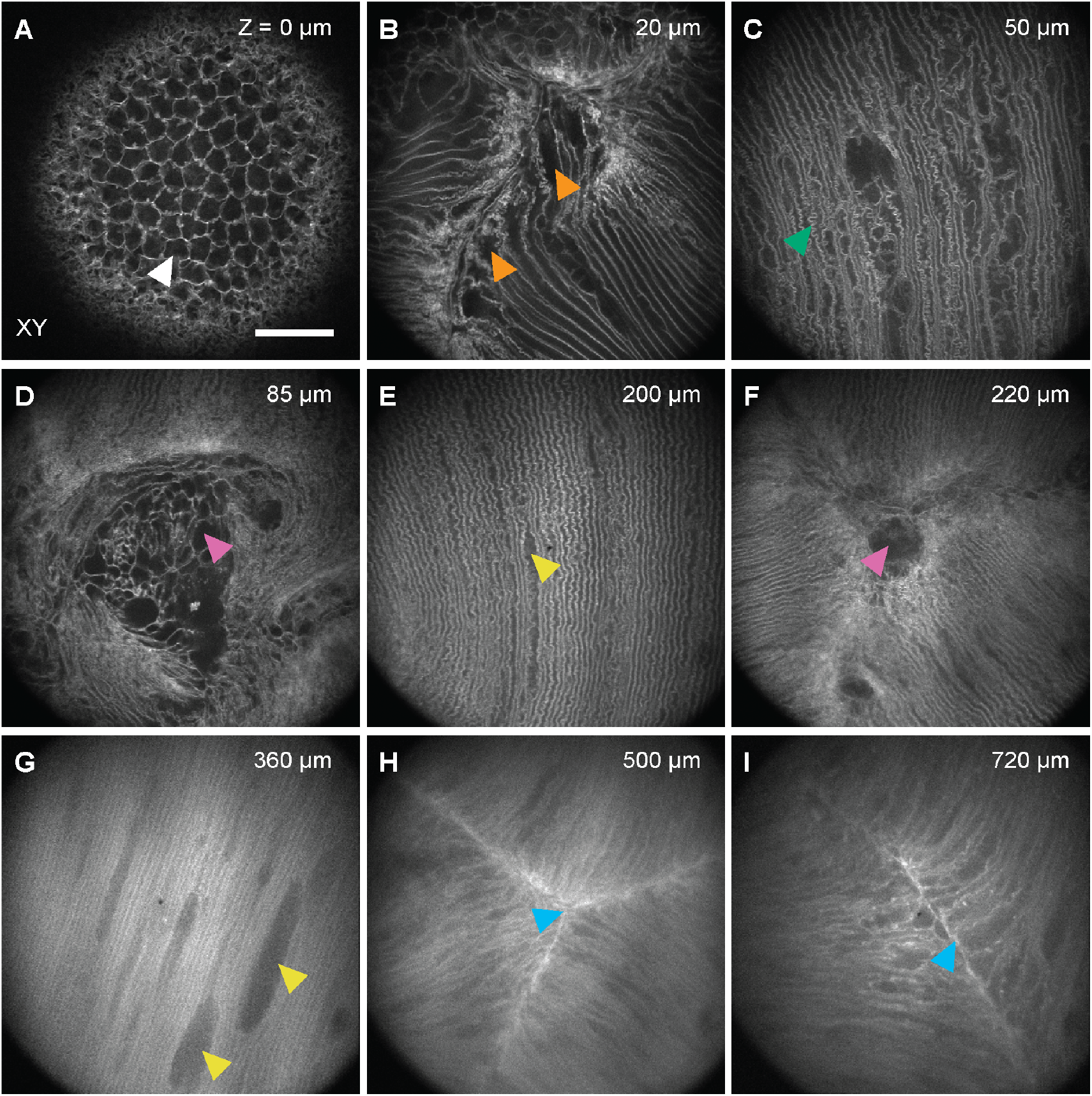
In vivo AO 2PFM visualizes features of epithelium and fiber cell organization in mouse lenses. (**A-I**) Representative images of cellular features at various depths visualized by AO 2PFM in the anterior lenses of the mice employed in the study. (**A**) Lens epithelium, (**B**) disorganized and broadened fiber cell ends, (**C**) non-uniform superficial fiber cells with excessive interdigitation, (**D, F**) anterior voids, (**E, G**) mostly uniform rows of fiber cells with enlarged vacuoles, (**H, I**) tight suture structures at depth. Depth noted in each panel is relative to the epithelium. Arrowheads point to features described in the main text. Scale bar: 50 μm.

In vivo, we observed structural organizations at the anterior part of the mouse lens that were consistent with the existing model of lens cellular organization. We found that the epithelium on the anterior surface of the lens capsule is characterized by connected polygonal cells (**Fig. 2A**). Below the epithelial cells, we found fiber cells at various depths in the central region of the anterior lens (**Fig. 2B-I**). When we imaged the lenses along the visual axis, we could also visualize the tight Y-shaped anterior sutures at the deeper regions of the anterior lenses (cyan arrowheads, **Fig. 2H-I**). In addition to these previously reported features of cellular organization in the lenses, we found some novel features in vivo. Unlike the ideal picture of identically shaped fiber cells forming highly ordered structures, we found irregularly-shaped fiber cells at shallow anterior regions (up to ~200 μm below epithelium) of the lens capsule (**Fig. 2B-C**). Depending on the orientation of the lens under the microscope objective and imaging depth, we found either fiber cell end broadening at the suture interface (orange arrowheads, **Fig. 2B**) or fiber cells with larger extent of interdigitation (green arrowhead, **Fig. 2C**) than that observed in the deeper regions. Next, we found large voids (pink arrowheads, **Fig. 2D,F**) with non-uniform fiber cell distribution at various depths along the anterior sutures of the lenses. These voids varied in size and shape along the depth of the lenses and across the animals imaged in this study. We also observed ellipsoidal enlarged vacuole-like structures within the lens fiber cells ranging from superficial to deeper regions of the lens (yellow arrowheads, **Fig. 2E,G**). In the following sections, we further investigated these features of fiber cell organization in vivo in 3D and tracked them longitudinally in growing mice.

### Anterior lens has void-like structures near suture

Prior imaging studies using sections of dissected mice lenses have established the presence of suture structures, where the tips of the elongating lens fiber cells meet under the apical surface of the epithelium and along the posterior capsule^25^. These sutures were described as Y-shaped compact interfaces between the fiber cell tips converging continuously along the visual axis. Given our surprising observation of anterior voids, we leveraged the 3D imaging capability of our AO 2PFM system to image approximately along the visual axis (**Fig. 3A**) to visualize the axial extension of the anterior voids near the lens suture. We performed AO measurements at 100 μm and 300 μm under the epithelium and applied the respective corrective wavefronts for imaging in the depth ranges of 0-200 μm and 200-400 μm (**Fig. 3B**) with an axial step size of 1 μm.

**Figure 3.**
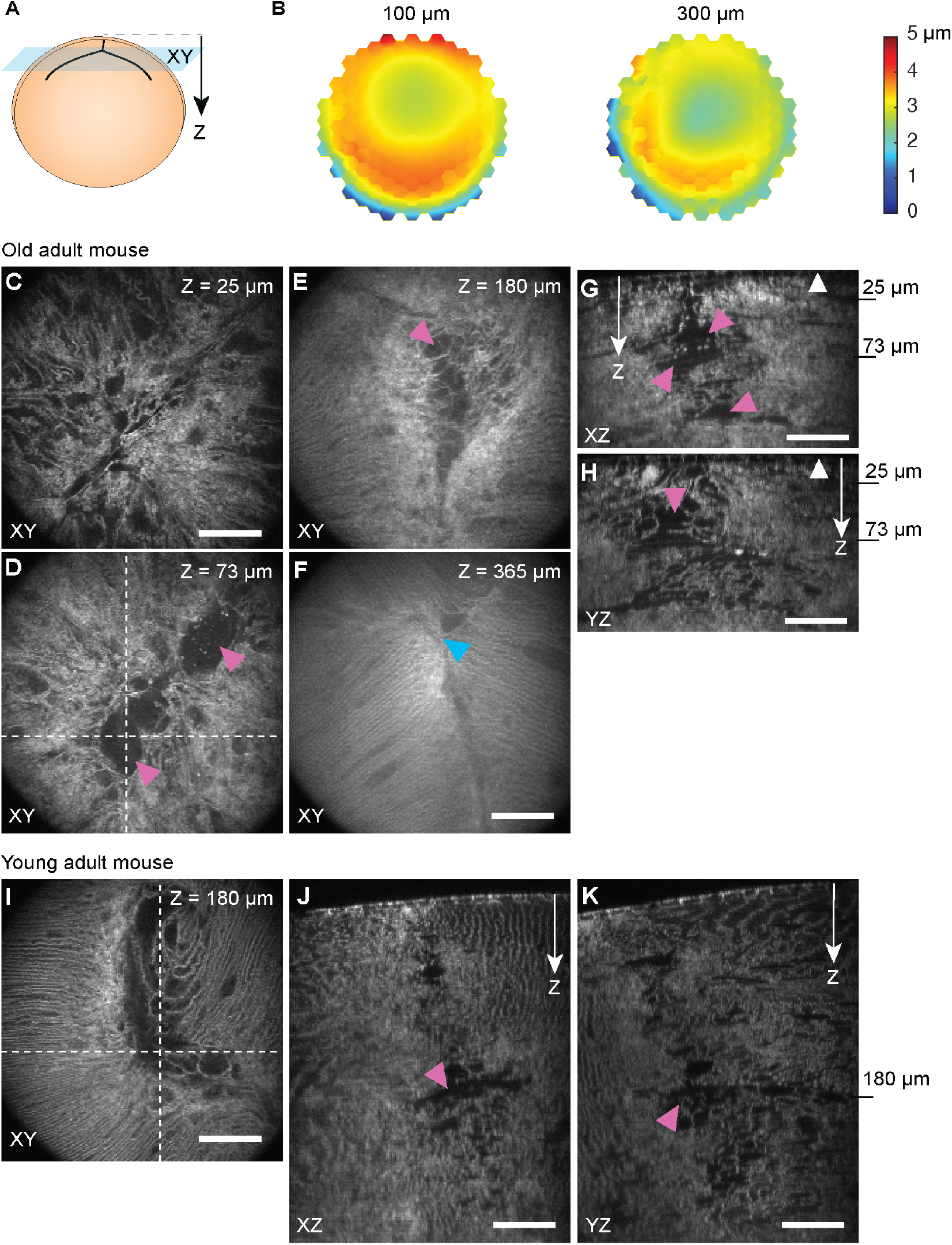
AO 2PFM enables 3D profiling of the anterior voids near suture in vivo. **(A)** The approximate orientation of the X, Y, and Z axes in the image stacks are shown with respect to the lens with anterior Y-suture at the top. (**B**) Corrective wavefronts acquired at 100 μm and 300 μm below epithelium. **(C-F)** Representative XY images of anterior voids in an older (35 weeks) adult mouse lens near anterior suture at different depths. (**G-H**) XZ and YZ cross-section of the anterior voids along the dashed lines in **D**, respectively. (**I-K**) Representative XY image, as well as XZ and YZ cross-sections (along the dashed lines in **I**) of a young (6 weeks) mouse lens. White arrowheads: location of epithelium; White arrows: direction along the imaging depth starting from the epithelium at the top. Scale bars: 50 μm.

When we probed the anterior lens of older adult mice (n = 6, 23-37 weeks), in vivo imaging revealed a lack of well-defined sutures in the shallow regions under the epithelium (**Fig. 3C-E**). Instead, we found that the anterior regions of the lens were marked by the presence of void-like structures that were either regions devoid of any cells or regions with sparse disorganized fiber cells (pink arrowheads, **Fig. 3D-E**). These voids (n = 6) had a variety of shapes and sizes across the anterior lenses with a mean maximum dimension of 128 μm and standard deviation of 37 μm across the older mice. At the deeper regions of the lens, however, we found relatively well-defined Y-shaped sutures with dense fiber cell packing (cyan arrowhead, **Fig. 3F**). The axial extension of these voids can be visualized by examining the XZ and YZ cross-sections of the anterior lens (**Fig. 3G-H**). While the anterior voids were not always contiguous through the lens, the non-uniform distribution of sparse fiber cells in them hints at a porous microenvironment along the anterior suture. We also examined younger mice (n = 7, 6 weeks) and observed that anterior voids were also present at young age (pink arrowheads, **Fig. 3I-K**). However, the voids (n = 7) were slightly smaller with a mean maximum dimension of 90 μm and standard deviation of 34 μm.

### Anterior lens fiber cells away from suture show depth-dependent organization and are marked by enlarged vacuole-like structures

We imaged away from the suture to avoid the anterior voids and captured fiber cell organization at multiple depths (axial step size of 2 μm, **Fig. 4A-P**) in the anterior lens approximately along the visual axis. We used the same AO approach as above to measure aberrations at 100 μm, 300 μm, and 500 μm under the epithelium and applying the obtained corrective wavefronts for imaging at nearby depths in young (n = 8, 8 weeks) and old (n = 5, 23-37 weeks) adult mice.

**Figure 4.**
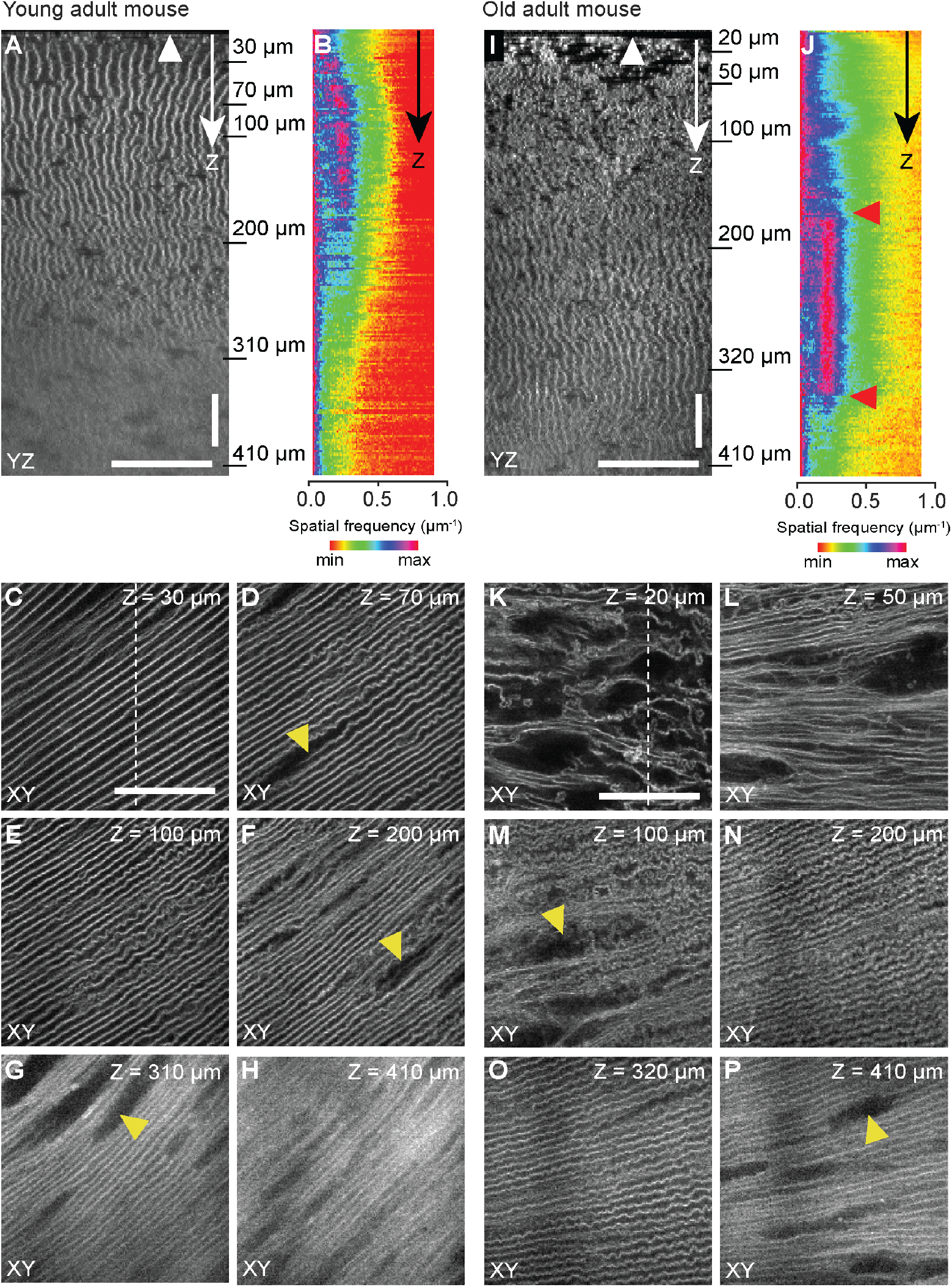
AO 2PFM enables structural characterizations of fiber cell organization away from suture across depths in vivo. (**A**) YZ cross-section of fiber cells in an example young mouse (8 weeks). (**B**) Radially averaged profiles of the Fourier transforms of XY images acquired at an axial step size of 2 μm. (**C-H**) Representative XY images at Z locations indicated in **A**. (Cross-section in **A** is visualized along the dashed line in **C**.) (**I**) YZ cross-section of fiber cells in an example older adult mice (34 weeks). (**J**) Radially averaged profiles of the Fourier transforms of XY images acquired at an axial step size of 2 μm. (**K-P**) Representative XY images at Z locations marked in **I**. (Cross-section in **I** is visualized along the dashed line in **K**.) White arrowheads: location of epithelium; White arrows: direction along the imaging depth starting from the epithelium at the top. Scale bars: 50 μm.

In the young mice, we observed neatly organized fiber cells right below the epithelium and a higher compaction of fiber cells in the deeper regions of the anterior lens (**Fig. 4A-H**). The differences in fiber cell organization with depth can be observed from both the YZ cross-section (**Fig. 4A**) and the depth-dependent radially averaged profiles of the Fourier transforms of the XY images (**Fig. 4B**), which indicate two depth ranges with differential organization. The superficial region that extended to ~250 μm below the epithelium had highly aligned and periodically arranged fiber cells, leading to strong high spatial frequency components in their XY images. Nevertheless, even within this region of high order, we found enlarged vacuole-like structures (yellow arrowheads, **Fig. 4D,F**). Below ~250 μm, although the packing of fiber cells remained compact, vacuoles became more pervasive (yellow arrowheads, **Fig. 4G**), which substantially disrupted the local order and suppressed the high spatial frequency components of the fiber cell images.

The organization of fiber cells in lenses of older adult mice (**Fig. 4I-P**) showed marked differences from that of young mice, with three distinct depth ranges as indicated by the YZ cross section and the radially averaged profiles of the Fourier transforms of the XY images (**Fig. 4I-J**). The fiber cells in the shallowest region (within ~150 μm under epithelium) were much more disorganized in their organization than those in the young mice (**Fig. 4K-M**), with the disorganization especially severe for the fiber cells immediately under the epithelium (**Fig. 4K**). Beyond the shallowest region, fiber cell organization became very ordered, forming well aligned rows with distinct peaks at higher spatial frequency (~180–350 μm, between red arrowheads, **Fig. 4J**). Comparing the XY images of fiber cells at similar depths (e.g., **Fig. 4E-G** versus **Fig. 4M-O**), we found that fiber cells in the adult mice exhibited a larger extent of interdigitation than those in the young mice. Beyond 350 μm, fiber cells became more compactly packed with more vacuoles (**Fig. 4P**), similar to the deeper cells in the young mice.

### In vivo 2PFM unveils large cavities in deeper regions of the lens

We further leveraged the ability of AO 2PFM for deep tissue imaging to investigate fiber cell organization deep into the lens core (600–980 μm, **Fig. 5**). At these larger depths, the pupil, even though dilated, started to block the marginal rays of the excitation light. As a result, the effective numeric aperture (NA) and consequently the spatial resolution of our 2PFM were reduced. The reduced NA also led to a more limited performance of AO correction^34^ (**Fig. S1**). As a result, only aberration correction for the optical system was applied. Nevertheless, the achievable resolution enabled us to observe large cavities in the deeper lens regions in both young (n = 4, 8 weeks; **Fig. 5A-C**) and older (n = 5, 34-37 weeks**; Fig. D-G**) mice. These cavities had well-defined borders and a complete absence of cells within them (pink arrowheads, **Fig. 5A-G**). We also saw appreciable accumulation of concentrated fluorophores in some of these cavities (cyan arrowheads, **Fig. 5E-G)**. The cavities in both young and older mice possessed a variety of shapes and sizes across the different mice within the same age group. The volumes of the cavities in the young mice ranged from 5.1 × 10^4^ μm^3^ to 1.7 × 10^6^ μm^3^ with an average volume and standard deviation of 6.2 × 10^5^ μm^3^ and 7.6 × 10^5^ μm^3^, respectively (n = 4). In the older mice, cavities were larger with volumes ranging from 1.2 × 10^5^ μm^3^ to 1.8 × 10^6^ μm^3^ with an average volume of 1.1 × 10^6^ μm^3^ and standard deviation of 8.6 × 10^5^ μm^3^ (n = 5).

**Figure 5.**
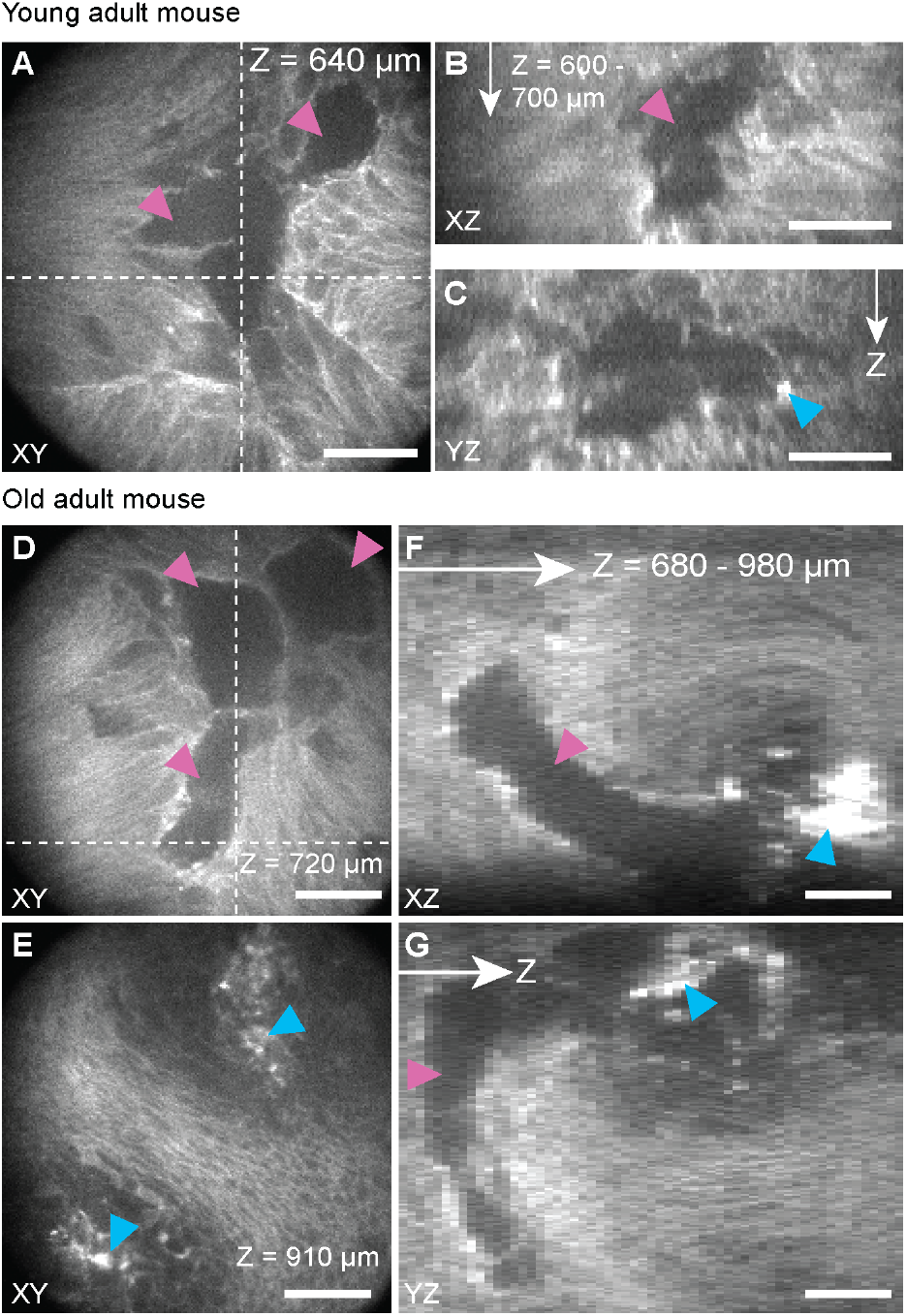
In vivo 2PFM imaging of large cavities in the lens core. (**A-C**) Representative XY, XZ, and YZ cross-sections (along the dashed lines in **A**) of a large cavity 600–700 μm beneath epithelium in a young mouse (8 weeks). (**D-G**) Representative XY images as well as XZ and YZ cross-sections (along the dashed lines in **D**) of a large cavity 680–980 μm beneath epithelium of an older mouse (34 weeks). White arrows in XZ and YZ cross-sections: axial depth direction. Scale bars: 50 μm.

### AO 2PFM enables longitudinal monitoring of lens growth in vivo

The success of non-invasive in vivo imaging enabled us to perform longitudinal monitoring of the dynamic changes in lens structure in the same animals during growth. We imaged the lenses of 5 mice from week 6 to week 12 (**Fig. 6**) every two weeks. Using anterior sutures and voids as landmarks, we were able to repeatedly locate and image the same volumes of lens tissue over weeks. Even though the incorporation of the newly elongated fiber cells from the equator resulted in additional layers of fiber cells over existing voids as visualized in the XZ and YZ cross-sections (orange line segments, **Fig. 6A-G**), the shapes and sizes of, as well as the distances between these voids remained stable with time (yellow line segments, **Fig. 6A-G**). The distance between the anterior void and the deeper cavity also remained nearly constant with time (yellow line segments, **Fig. 6H-I**), despite the continuous growth during this period as evinced by their displacement compared to the epithelium. Together, these results show that AO 2PFM enables longitudinal localization and repeated measurements of morphological features of the lenses in vivo.

**Figure 6.**
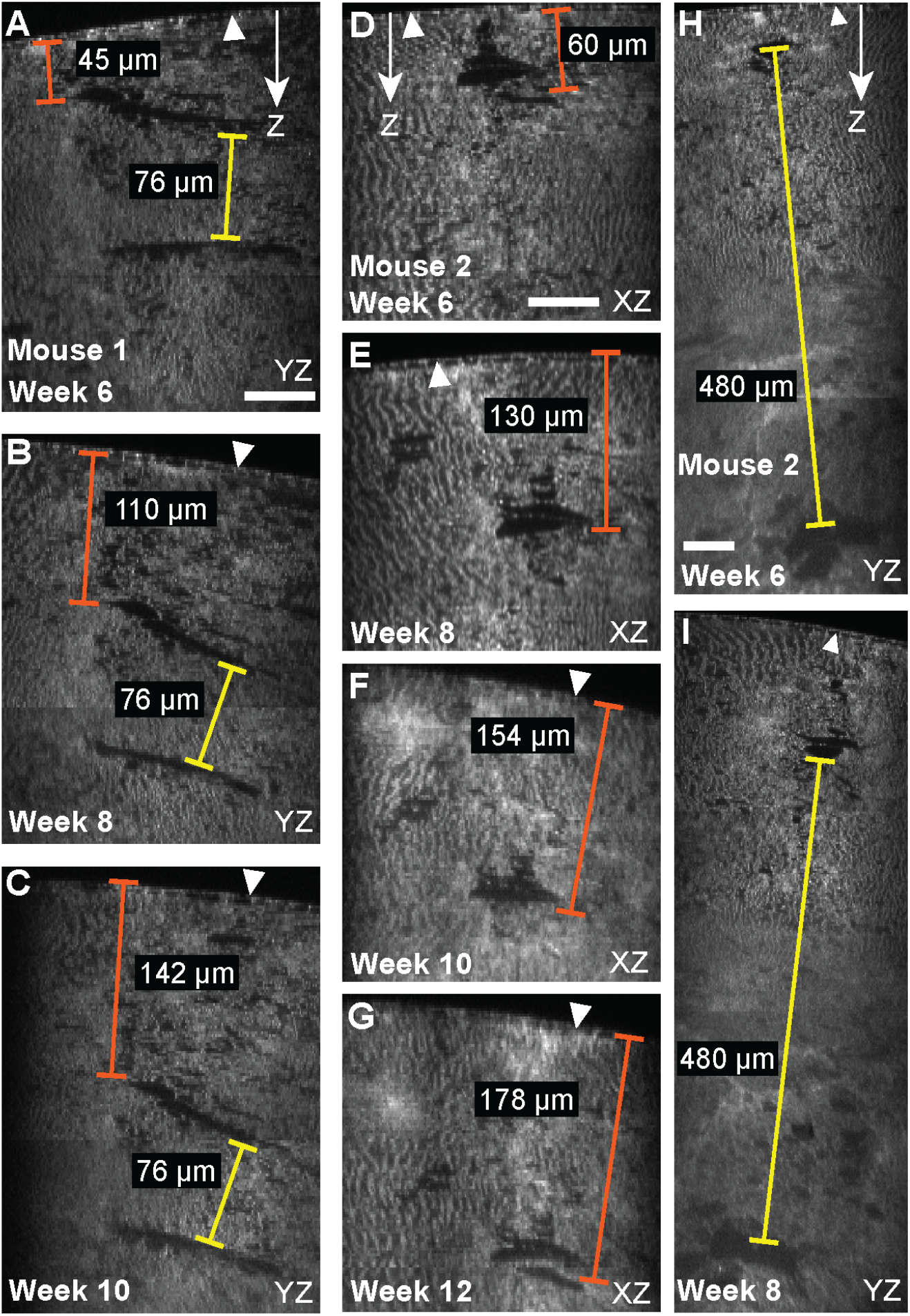
Longitudinal monitoring of lens growth in vivo using AO 2PFM. (**A-C**) Representative YZ cross-sections of the same lens (Mouse 1) including the same anterior voids imaged at weeks 6, 8, and 10. (**D-G**) XZ cross-sections of another mouse (Mouse 2) with the same anterior voids imaged at weeks 6, 8, 10, and 12. (**H-I**) YZ cross-sections of Mouse 2 including a shallow anterior void and a deep cavity imaged at weeks 6 and 8. White arrowheads: location of epithelium; White arrows: direction along the imaging depth starting from the epithelium at the top. Scale bars: 50μm.

## Discussion

Ocular lens is of considerable interest to biology due to both its importance to vision and its continuous growth over the lifespan of the organism^40^. Prior research has attempted in vivo microscopic examination of lens development in zebrafish, a simpler vertebrate model^41–44^. However, the lens formation and growth in mammals are different in size and cell growth rates^45^. Therefore, for understanding the organization of lens cells during postnatal development and aging, it is crucial to be able to perform noninvasive in vivo microscopic imaging of lenses in living mice. We chose to image mouse strains with tdTomato fluorescent protein expression in the plasma membrane of the lens cells, because they have uniform labeling across the lens fibers from the periphery to the core^26^. The brightness and photostability of tdTomato and its 2P excitation in the near infrared (NIR) region make it an ideal candidate for in vivo imaging using AO 2PFM, which requires bright structures for sensing the wavefront distortion caused by sample-induced aberrations^37^. The resolution improvement after correcting these aberrations enabled us to investigate structural features such as voids and vacuoles and the detailed arrangement of fiber cells at subcellular level. The optical sectioning capability and large penetration depth of 2PFM allowed us to image ~1 mm deep into the core of the mouse lens, revealing the existence of large cavities towards the center of the lens.

Previous in vitro studies of enucleated lenses used confocal fluorescence microscopy and electron microscopy to characterize lens cellular organization^20–24^ and established the molecular and structural mechanisms of lens growth, such as the loss of organelles by the proliferating cells at the equator and the incorporation of layers of fiber cells into the core of the lens as they age^1^. With in vivo AO 2PFM measurements, we observed similar features such as the arrangement of epithelium and fiber cells in the anterior part of the lens, the differential fiber cell density across the depth, and their hexagonal cross section. However, a more complex picture of fiber cell organization emerges from our in vivo imaging data. Prior investigations on the anterior suture – using both confocal and electron microscopy – in enucleated vertebrate lenses described them as tight branched structures along the visual axis^25,46^.

Our in vivo lens imaging results from all the mouse strains tested revealed that the anterior sutures in the superficial regions of the lens are marked by complex voids with non-uniform fiber cell distribution and varying degree of openness. These anterior voids have not been reported in previous ex vivo studies and their exact composition remains to be determined. Such voids may collapse in enucleated lenses due to the absence of attached zonule structures and/or lens active flow transport system. It is unknown whether those voids are related to normal physiological function of the lens or result from cellular pathologic events during development. Their occurrence in both young and older mice suggests that they are probably not a feature of age-related dysregulation of cellular organization and may be a normal and critical part for lens growth and survival. We speculate that these voids may serve as reservoirs of nutrients and small molecules and are part of the inward flow pathway of lens circulation to facilitate a dynamic transport between the core of the avascular lens and the anterior chamber^3^. The observation of these anterior voids opens a new route of inquiries into the molecular transport between the lens periphery and the lens core from the exterior anterior and posterior chambers. These voids occur much less frequently in the deeper regions towards the lens core, where we found uniform Y-shaped sutures with tight and orderly fiber cell arrangement known to be required for the light transmission and formation of a focused image on the retina.

Away from the sutures, our observations of differential lens fiber morphology across depth are largely consistent with prior reports with higher compaction of fiber cells in the primary fibers at the core of the lens than the peripheral fiber cells^47,48^. Prior microtome section studies of human lenses focused on the fiber cell morphology in the equatorial plane to visualize and demarcate the epithelium, differentiating fiber cells, remodeling zone of disordered assembly of nucleated fiber cell, and transition zone composed of anucleated fiber cells arranged in an irregular pattern that precedes the compaction of anucleated fiber cells of the adult nucleus in human lenses^47,49^. While mouse lenses have been shown to recapitulate some aspects of aging in human lenses, they are known to be challenging for fixation and vibratome sectioning for ex vivo studies. In vivo imaging using 2PFM enables us to examine the cellular organization just under the epithelium in the lenses of older mice along the visual axis. Based on the distances from the lens surface, we believe that the disorganized fiber cells observed in the superficial depths below epithelium likely belong to the elongated fiber cells from the remodeling zone and early transition zone to the anterior suture region. At larger depths, we saw similar compaction of fiber cells along the visual axis to that observed along the equatorial plane in these and other previous studies^47–49^. Our findings of distinct zones of cellular organization of fiber cells in the younger and older mice along with the observation of ordered fiber cells under epithelium in younger mice agree with the prior electron microscopy characterization of morphological degradation of fiber cells with age^50^.

We observed enlarged vacuoles throughout the anterior lenses of both younger and older adult mice. We hypothesize that these vacuoles may be pockets filled with fluids that are transported in and out of the lens core for their nutrition and maintenance. Similar vacuoles have been observed in previous investigations of lens sections using confocal and electron microscopy. Shi et al. observed similar structures in the fiber cells that were proposed to be a result of Lim2-dependent cellular fusions^51^. Baba et al. reported similar vacuoles in Sall1-gfp mice that became more pronounced with the knockdown of Sall1^52^. Their high-magnification electron microscopy data showed that these vacuoles are extracellular spaces between the fiber cells. While not explicitly discussed, narrow gaps between the fiber cells similar to the enlarged vacuoles observed in our data could be seen in the confocal microscopy images of fiber cells in tdTomato mice in Parreno et al^25^. Imaging fiber cells deep inside the anterior lens in vivo, our approach enabled us to observe the distribution of these vacuoles in 3D and find that these vacuoles to be more prevalent at larger depths. Additionally, we were able to observe large cavities in the deeper regions of the anterior lens, which were not reported previously. We speculate that these cavities may reflect cellular alterations due to membrane/protein segregation and accumulation. Furthermore, their presence in the younger lenses and temporal stability indicate that they may be resultant of irreversible degradation of lens cells.

In summary, our results show that AO 2PFM enables in vivo imaging of the anterior lens morphology in the mouse eye by leveraging mouse models that express fluorescence in the cell membrane of lens cells. Correcting sample-induced aberrations with AO, we can maintain subcellular resolution and image contrast deep inside the lens. In vivo imaging of the intact lens, with its active transport and support structures in place, revealed novel features of fiber cell organization in the lens including suture-associated superficial anterior voids, vacuoles, and large cavities at depth. Our ability of longitudinal monitoring of the same anterior features during lens growth is expected to facilitate further biological investigations into changes in lens cellular organization, such as tracking the development of presbyopia and formation of cataract due to aging and pathological transformation in advanced genetic models. The subcellular resolution of our method would also allow the lens biology field to monitor how lens cells respond to therapeutical manipulations with cellular and molecular details.

## Methods

### Animal models

The animal experimental protocols were approved by the Institutional Animal Care and Use Committee at the University of California, Berkeley. Experiments followed the National Institutes of Health guidelines for animal research. We employed mice expressing membrane-targeted tdTomato (strain 007676, The Jackson Laboratory) in the C57BL/6 background to serve as wildtype (WT) mice (n = 6) for these experiments^26^. The klotho-related protein KLPH (lctl) knockout mice was provided by Dr. Melinda Duncan at University of Delaware. This mouse strain was originally generated by Dr. Graeme Wistow at National Eye Institute^21^. We also employed KLPH-KO (n = 7) and KLPH-Het (n = 5) double transgenic mice that were generated by mating the KLPH knockout mice with transgenic mice expressing membrane targeted tdTomato (strain 007676, The Jackson Laboratory). All mice were maintained at the C57BL/6J staining backgrounds in the lab. Animals were housed with free access to food and water and exposed to 12h:12h light:dark cycle. All experiments were performed in live mice between the ages of 6 weeks and 9 months. The genotype, sex and age at imaging have been tabulated in **Table ST1** (**Supporting Information**).

### Animal preparation for AO 2PFM imaging

In vivo imaging experiments were performed under isoflurane anesthesia (~1.5% by volume in O_2_). The anesthetized mice were placed on their side and mounted on a bite bar attached to a dual-axis goniometer stage (Thorlabs, GNL18) with two rotational degrees of freedom and equipped with a nose cone for continuous gas anesthesia during imaging. The body temperature of the mouse was maintained with a heating pad. The pupil of the anesthetized mice was dilated by the application of one drop each of 2.5% phenylephrine hydrochloride (Paragon BioTeck, Inc) and 1% tropicamide (Akorn, Inc). To prevent corneal drying during the imaging session, we used a drop of eye gel (Genteal) between the dilated eye and a cover glass stably mounted on a holder. The animal was placed under the microscope objective by sandwiching a drop of water (immersion media) between the objective and the cover glass. The fine adjustment of the position of mouse eye with respect to objective was controlled using a motorized translational stage (Newport, Series 562) on which the mouse was placed. Since the suture is not aligned centrally to the pupil of the animal, we imaged slightly away from the center of the pupil to capture the features around the anterior suture.

### In vivo AO 2PFM imaging

In vivo imaging experiments were performed on a custom-built 2PFM system with direct-wavefront-sensing AO. The optical setup has been described in detail previously^35^. Briefly, the system consisted of a femtosecond Ti:Sapphire laser (Coherent, Chameleon Ultra II) tuned to 1000 nm, and its output beam scanned with a pair of optically conjugated galvanometer mirrors (Cambridge Technology, 6215H). The beam was relayed from the galvos to a segmented deformable mirror (DM, Iris AO, PTT489) using a pair of achromatic lenses (Edmund Optics, 49-359-INK and 49-368-INK) and the DM was conjugated to the back pupil plane of the objective (Nikon, CFI Apo LWD 25x, 1.1NA and 2-mm working distance) using another pair of achromatic lenses (Newport, PAC090 and Edmund Optics, 49-363-INK). The objective was mounted on a piezoelectric stage (Physik Instrumente, P-725.4CD PIFOC) for controlling its axial motion. The correction collar on the objective was adjusted to compensate for the spherical aberration due to the cover glass on top of the mouse eye. The fluorescence signal from the ocular lens was reflected by a dichroic mirror (Semrock, Di02-R785-25×36), filtered (Semrock, FF01-607/70-25), and detected by a photomultiplier tube (PMT, Hamamatsu, H7422-40). Direct wavefront sensing for AO was achieved by moving the dichroic mirror out of the path and descanning the emitted fluorescence by the galvo pair to a Shack-Hartmann (SH) sensor comprised of a lenslet array (Advanced Microoptic System GmbH, APH-Q-P500-R21.1) and a camera (Hamamatsu, Orca Flash 4.0). We scanned an area of 19 μm × 19 μm for ~10 s to capture the SH pattern for wavefront sensing. The aberrated wavefront was reconstructed using the measured shifts of the SH pattern foci and a custom MATLAB code. The corresponding corrective pattern was applied to the DM for achieving aberration-free imaging.

To accurately assess the AO improvement due to the correction of sample-induced aberrations, we used a modal-based optimization approach to correct for the system aberrations due to the imperfections and misalignment of the microscope optics, prior to making the in vivo measurements in the ocular lenses^53^. Here, we imaged fluorescence beads and applied 21 values (−0.2 ~ 0.2 μm rms at an increment of 0.02) for each of the first 21 Zernike modes excluding piston, tip, tilt, and defocus. For each Zernike mode the optimal value was determined by maximizing the fluorescence intensity of the sample and it was applied to the DM before proceeding to the next Zernike mode. We used the SH pattern obtained after correcting the system aberrations as the SH reference for calculating sample-induced aberrations. All the images labeled as “No AO” in the manuscript were taken after performing the system correction, which represented the best performance of the microscope without correcting for the sample-induced aberrations. The laser power and image acquisition parameters for all the experiments are provided in **Table ST1** (**Supporting Information**).

### Image processing and analysis

All image processing and analysis have been performed using ImageJ/FIJI (NIH)^54^, Python scripts, and MATLAB (Mathworks) software. 2D image registration using TurboReg plugin was performed to remove motion-induced artifacts. For 3D image stacks, we normalized the individual 2D XY images to have the same means to compensate for the loss of signal with depth and better illustrate the features in the deeper regions in the orthogonal cross sections. For calculating the spatial frequency space representations (i.e., the Fourier transforms) of the XY images, we applied a Gaussian Blur filter in ImageJ with a blur radius of 1 pixel to these images to reduce pixelation artifacts. Logarithmic scale was used to show the frequency space representations. The volumes of the deeper cavities were estimated using the equation V = (*π*/6) x (length) x (width) x (height).

## Acknowledgements

X.G., C.H.X., and S.K.P. acknowledge the support from the National Institutes of Health grants (R01EY013849; R01EY031253; P30EY003176). Q.Z., Y.Y., and N.J. acknowledge the support from the National Institutes of Health grant (U01NS118300).

