## Supporting Information for "Adaptive optical two-photon fluorescence microscopy probes cellular organization of ocular lenses in vivo"

The authors disclose no potential conflicts of interest.

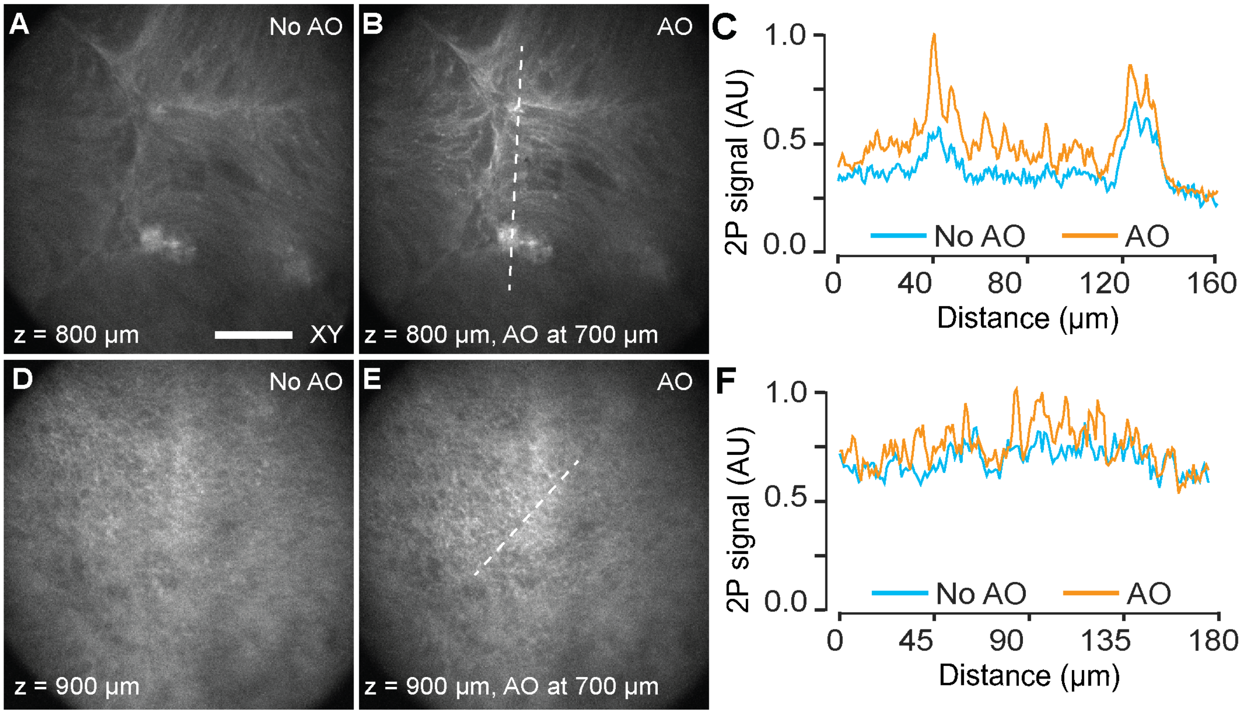


**Figure S1. In vivo AO 2PFM in the deeper regions of anterior lens.** (**A-B**) XY images of fiber cells at 800 μm below epithelium (**A**) without and (**B**) with AO correction at a depth of 700 μm. (**C**) Signal profiles along the dashed line in **A** and **B**. (**D-E**) XY images of fiber cells at 900 μm below epithelium (**D**) without and (**E**) with AO correction at a depth of 700 μm. (**F**) Signal profiles along the dashed line in **D** and **E**. Scale bar: 50 μm.

### Table ST1. Animal characteristics and imaging parameters for all the figures.

| Parameters | Fig. 1 | | | | Fig. 2 | | | | | | | | |
|---|---|---|---|---|---|---|---|---|---|---|---|---|---|
| Panel | A, B | D, E | J, K | M, N | A | D | F | G | B | H | C | E | I |
| Mouse line | KLPH-KO | | WT | | KLPH-Het | | | | KLPH-KO | | WT | | |
| Age (weeks) | 12 | | 31 | | 16 | 21 | 21 | 16 | 12 | 11 | 30 | | |
| Sex | F | | | | F | | | | | | | | |
| Illum. power (mW) | 3 | | 5.5 | | 2 - 10 | | | | | | | | |
| FOV(µm x µm) | 228 x 228 | 76 x 76 | 228 x 228 | 76 x 76 | 228 x 228 | | | | | | | | |
| Pixel size (µm) | 0.57 | 0.19 | 0.57 | 0.19 | 0.57 | | | | | | | | |
| Axial step (µm) | / | | | | / | | | | | | | | |
| Stack thickness (µm) | / | | | | / | | | | | | | | |
| XY Integration time (s) | 20 | | 20 | | 20 | | | | | | | | |

| Parameters | Fig. 3 | | | | Fig. 4 | | | | Fig. 5 | | | |
|---|---|---|---|---|---|---|---|---|---|---|---|---|
| Panel | C-F | G-H | I | J-K | A | C-H | J | K-P | A | B-C | D-E | F-G |
| Mouse line | WT | | WT | | WT | | KLPH-Het | | WT | | KLPH-KO | |
| Age (weeks) | 35 | | 6 | | 8 | | 34 | | 8 | | 34 | |
| Sex | F | | M | | F | | F | | F | | F | |
| Illum. power (mW) | 2 - 7.5 | | 3.3 - 7.5 | | 3.2 - 8.5 | | 1.1 - 3.7 | | 6.5 | | 7.4 | |
| FOV(µm x µm) | 228 x 228 | 228x144 | 228x228 | 228x294 | 114 x 420 | 114 x 114 | 114 x 420 | 114 x 114 | 228x228 | 228x98 | 228 x 228 | 228x295 |
| Pixel size (µm) | 1.14 | | | | 0.57 | | | | 1.14 | | 1.14 | |
| Axial step (µm) | 1 | | | | 1 | | | | 2 | | 5 | |
| Stack thickness (µm) | / | 144 | / | 294 | 420 | / | 420 | / | / | 98 | / | 295 |
| XY Integration time (s) | 5 | | | | 5 | | | | 5 | | 5 | |

| Parameters | Fig. 6 | | | | | | | | | Fig. S1 |
|---|---|---|---|---|---|---|---|---|---|---|
| Panel | A | B | C | D | E | F | G | H | I | A-B, D-E |
| Mouse line | WT | | | | | | | | | WT |
| Age (weeks) | 6 | 8 | 10 | 6 | 8 | 10 | 12 | 6 | 8 | 31 |
| Sex | M | M | M | F | F | F | F | F | F | F |
| Illum. power (mW) | 2 - 10 | | | | | | | | | 5.5 |
| FOV(µm x µm) | 228 x variable depth | | | | | | | | | 228 x 228 |
| Pixel size (µm) | 1.14 | | | | | | | | | 0.57 |
| Axial step (µm) | 2 | | | | | | | | | / |
| Stack thickness (µm) | 294 | 294 | 320 | 228 | 228 | 228 | 228 | 578 | 646 | / |
| XY Integration time (s) | 5 | | | | | | | | | 20 |
